# Battery-Free Optoelectronic Patch for Photodynamic and Light Therapies in Treating Bacteria-Infected Wounds

**DOI:** 10.1101/2023.12.13.571417

**Authors:** Zhao Xue, Wenxin Chou, Yixuan Xu, ZiYi Cheng, Xuechun Ren, Tianzhen Sun, Wenbin Tong, Yang Xie, Junyu Chen, Nuohan Zhang, Xing Sheng, Yongtian Wang, Hongyou Zhao, Jian Yang, He Ding

## Abstract

Light therapy is an effective approach for the treatment of a variety of challenging dermatological conditions. In contrast to existing methods involving high doses and large areas of illumination, alternative strategies based on wearable designs that utilize a low light dose over an extended period provide a precise and convenient treatment. In this study, we present a battery-free, skin-integrated optoelectronic patch that incorporates a coil-powered circuit, an array of microscale violet and red light emitting diodes (LEDs), and polymer microneedles (MNs) loaded with 5-aminolevulinic acid (5-ALA). This polymer MNs, based on the biodegradable composite materials of polyvinyl alcohol (PVA) and hyaluronic acid (HA), serves as light waveguides for optical access and a medium for drug release into deeper skin layers. Unlike conventional clinical photomedical appliances with a rigid and fixed light source, this flexible design allows for a conformable light source that can be applied directly to the skin. In animal models with bacterial-infected wounds, the experimental group with the combination treatment of metronomic photodynamic and light therapies reduced 2.48 log_10_ CFU mL^−1^ in bactericidal level compared to the control group, indicating an effective anti-infective response. Furthermore, post-treatment analysis revealed the activation of proregenerative genes in monocyte and macrophage cell populations, suggesting enhanced tissue regeneration, neovascularization, and dermal recovery. Overall, this optoelectronic patch design broadens the scope for targeting deep skin lesions, and provides an alternative with the functionality of standard clinical light therapy methods.

## Introduction

Healthy skin is a robust barrier, protecting internal tissues from pathogens, ultraviolet rays, and other external hazards(*1*). Impaired healing of wounds, however, resulting from severe burns, traumatic injuries, surgical procedures, or chronic ailments such as diabetes, represents a considerable clinical issue. Such conditions often lead to a diminished quality of life, significant healthcare expenses, and an increased mortality risk (*2*). Meanwhile, the open, moist, and weeping wounds are conducive to bacterial colonization, further impeding healing by fostering an environment that aggravates the underlying conditions. The accumulation of bacteria and their toxins prompts an excessive release of inflammatory mediators, perpetuating an extended inflammatory phase that can derail the natural wound repair mechanisms(*3, 4*). Hence, interventions that inhibit bacterial colonization or infection and mitigate inflammation are pivotal for enhancing wound healing.

Light therapies have emerged as promising methods for treating various skin-related issues, including promoting wound closure and stimulating hair growth(*2, 5*).In addition, photodynamic therapy (PDT) combines a photosensitizer, excitation light, and oxygen to generate cytotoxic reactive oxygen species to efficiently destroy abnormal tissue(*6–11*). These therapies typically employ light in the visible spectrum. However, the penetration of light in this wavelength range into biological tissues is hindered by significant absorption and scattering, posing challenges for treating deep dermal conditions. Using implantable optical waveguides to deliver optical signals and energy directly into deep tissues can mitigate this issue. However, the rigid design and stiffness mismatch may induce negative effects such as tissue damage, inflammation, and rejection. Flexible, stretchable, and biodegradable polymeric materials, such as synthetic polymers, hydrogels, and bioderived materials, have been explored in both *in vitro* and *in vivo* studies to overcome these challenges(*12–18*). Polymer microneedles (MNs) made from biocompatible and biodegradable materials like hyaluronic acid (HA) and polyvinyl alcohol (PVA) provide a minimally invasive means of transdermal drug delivery and light transmission to the deeper layers of the skin(*19–23*). In addition, light treatment is currently confined mainly to specialized healthcare settings due to the impracticality and high cost of home-based devices for widespread use. The broad exposure area of such devices necessitates protective measures for unaffected skin and eyes, which complicates the treatment process. To overcome the limitations of such traditional light treatment setups, emerging techniques with wireless, wearable, and implantable systems with optoelectronics devices and related circuits have been developed(*23–30*). These small, lightweight devices provide a user-friendly and efficient alternative for administering low-light doses and facilitating long-term treatments.

This paper presents a battery-free optoelectronic patch system designed for light therapy application on infected traumatic skin wounds, which mainly incorporates coiled-powered LEDs and 5-ALA-infused biodegradable MNs. By embedding a photosensitizer within an optimal blend of water-soluble PVA and HA, these MNs act as both a medium for drug delivery and optical waveguides, delivering the drug and light into the deeper tissues. Numerical simulations and experimental data reveal that such a design can effectively increase the light transmission dose of deep skin. Effectively, this optoelectronic patch is capable of treating and resolving wound infections, especially beneficial for injuries prone to abrasions or cuts, provided it is applied promptly and correctly. Administered in a timely and suitable manner, this patch has the potential to successfully treat wound infections in mice through a combination of metronomic photodynamic therapy (mPDT) and light therapy. This proposed smart optoelectronic MNs patch may revolutionize the treatment of various complex skin conditions, offering a non-invasive, effective, and portable option for patient management.

## Results

### System Overview

Figures 1a depict the exploded view of wireless optoelectronic wound patch composed of multiple layers and components, mainly including drug-loaded biodegradable MNs, a UV–visible transparent epoxy encapsulation forming a bilayer, and multiple wavelength illumination with a violet LED and four red LEDs affixed to a copper-clad polyimide (PI) film that is approximately 100 μm thick (AP8535R, Pyralux Dupont). The flexible printed circuit board (FPCB) is fashioned into a thin, pliable layout through the use of a laser cutting tool (ProtoLaser U4, LPKF), resulting in conductive traces that enable the board to easily conform to the contours of a wound when attached with polyurethane (PU) medical tape. The FPCB integrates an antenna coil designed to resonate at a frequency of 13.56 MHz, allowing the harvesting of radio-frequency (RF) energy to illuminate the LEDs for light therapy. These devices and circuits are sequentially encapsulated with parylene (∼10 μm) and polydimethylsiloxane (PDMS, ∼100 μm) to provide electrical insulation and act as an adhesive for the MNs layer. As depicted in Figure 1b, beneath the protective layer is the replaceable drug-loaded MNs layer, which directly targets infected tissue for effective treatment. Furthermore, the LEDs in the patch remain functional even when the FPCB is flexed to a curvature radius as small as 9 mm, demonstrating impressive mechanical durability. With an approximate diameter of 18 mm, a mass of around 40 mg (Figure S1), and desirable mechanical flexibility, this smart patch emerges as an innovative and versatile solution for wireless light therapy applications.

**Figure 1.**
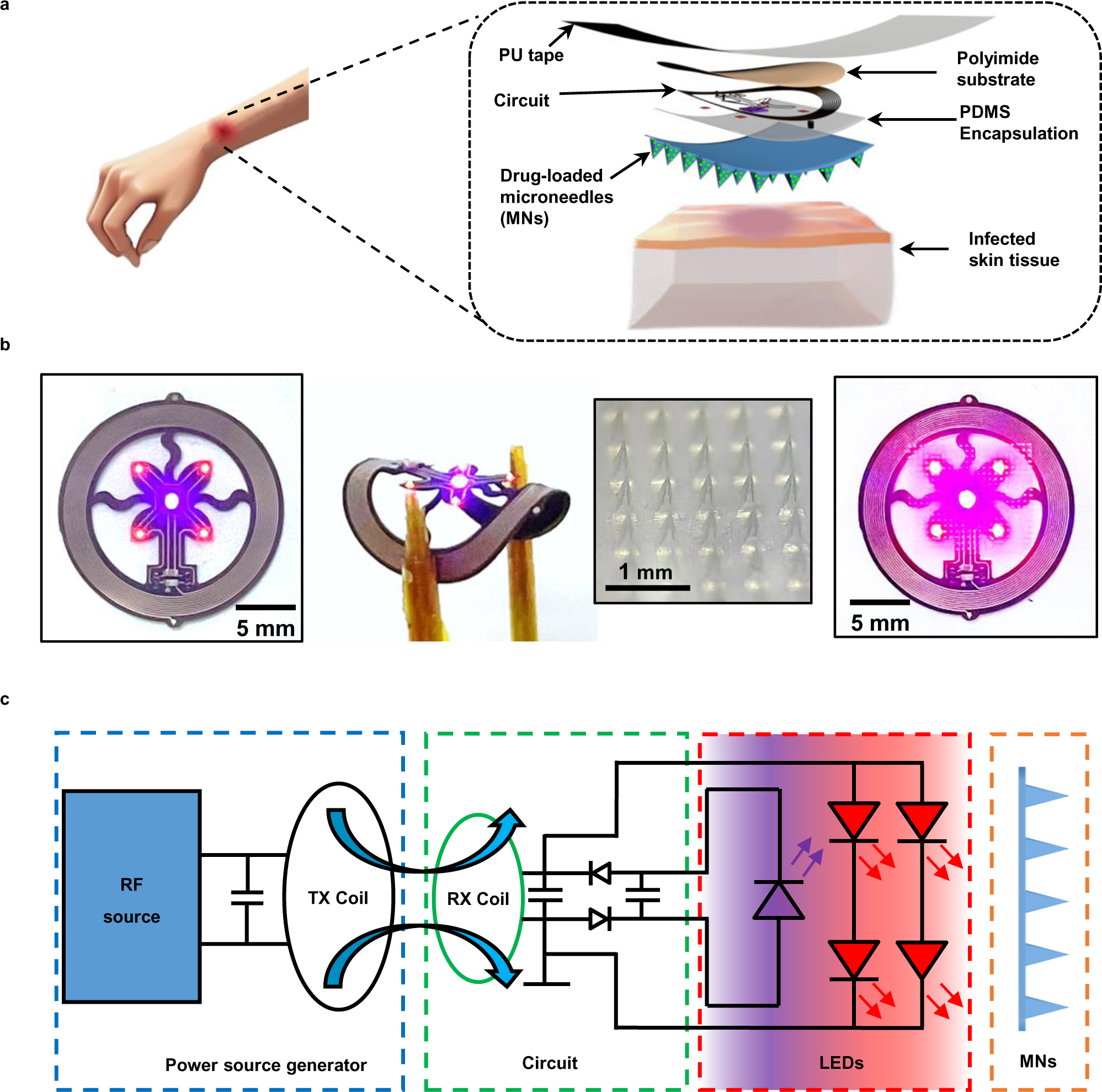
Battery-free optoelectronic patch for light therapy. (a) Schematic diagram placement and exploded view of the optoelectronic patch concept, primarily including microneedles (MNs) array layer and an encapsulated coil-powered LEDs system on a polyimide (PI) substrate, assembly by PU tape and covered to a wound on the skin. (b) Images of the illuminating flexible optoelectronic patch with four red LEDs and a violet LED in a flexible open architecture circuit powered by near-field magnetic resonant coupling, and integrated with a thin film replaceable MNs layer. (c) System block diagram outlining the overall system components, including a power source generator equipped with a radio-frequency (RF) source and transmitter coil (TX coil), and the optoelectronic patch includes wireless power receiving circuits with a receiver coil (RX coil), violet and red microscale LEDs, and MNs.

### Wireless Circuit Design

Figure 1c illustrates the structural design of the wireless optoelectronic device, including the antenna coil, LEDs, and relevant components. The antenna is a 10-turn circular coil with a diameter of 18 mm, and each coil has dimensions of 100 μm in width, 18 μm in thickness, and an inter-coil spacing of 80 μm. The chosen surface-mounted chips provide the coil with an optimal inductance of 4.6 μH and a resonant capacitance of 30 pF. This circuit design achieves resonance at the 13.56 MHz frequency through RF energy harvesting. The circuit features two types of LEDs: a violet LED with an emission wavelength of 405 nm in size of 200 μm × 400 μm × 50 μm and another red LED at an emission wavelength of 630 nm with size of 100 μm × 200 μm × 50 μm. In a specific arrangement, a single violet LED is encircled by four red LEDs, each spaced 3 mm apart from the other. The red LEDs are configured with two in series and two in parallel, a setup chosen because the activation voltage of two series-connected red LEDs is comparable to that of a single violet LED, as detailed in Figure S2. Furthermore, the violet LEDs operate as half-wave rectifiers. This design enables it to convert the incoming RF alternating current (AC) signals into direct current (DC) signals.

Figure 2a is the emission spectra of violet and red LEDs in the optoelectronic patch, in which the wavelengths of 405 nm and 630 nm align closely with the absorption peak of the loaded photosensitizer of 5-ALA induced protoporphyrin IX (PpIX) to ensure maximum photodynamic activation. An external four-loop antenna encircles a 20 cm × 15 cm cage and connects to an RF generator with an adjustable output power of 0–10 W and a frequency range of 0–100 Hz. When the RF generator operates at a power of 8 W, the emission intensities of the violet and red LEDs in the optoelectronic patch are 3 mW and 1.8 mW, respectively. Figures 2b, 2c, and S3 illustrate the light intensity distribution of the optoelectronic patch positioned in a horizontal plane of 3 cm and centrally positioned at a distance of 3–8 cm, respectively. Despite slight fluctuations in light intensity within the cage, the emission intensity of the LEDs is meticulously calibrated to guarantee efficacious light treatment throughout the areas of animal activity. Figure 2d indicates the coupled power by the coil correlates linearly to the external RF generator, a crucial factor for ensuring reliable illumination of the LEDs. In order to confirm the consistent performance of the LEDs for extended light therapy sessions, the stability of the light emission intensity over time and under different RF power levels has been characterized, as shown in Figure S4. Additionally, since the antenna coil is attached to the back of the animal, it may undergo fluctuations in power conversion due to body movements, thus affecting the LED emission intensities. Figure 2e highlights the trend of diminishing emission intensity as the angle of the patch increases from its initial flat position. Nevertheless, even with a bend or rotation of 60°, the intensity of the emitted light remains at 50% of its original strength, demonstrating the resilience of the system under practical conditions.

**Figure 2.**
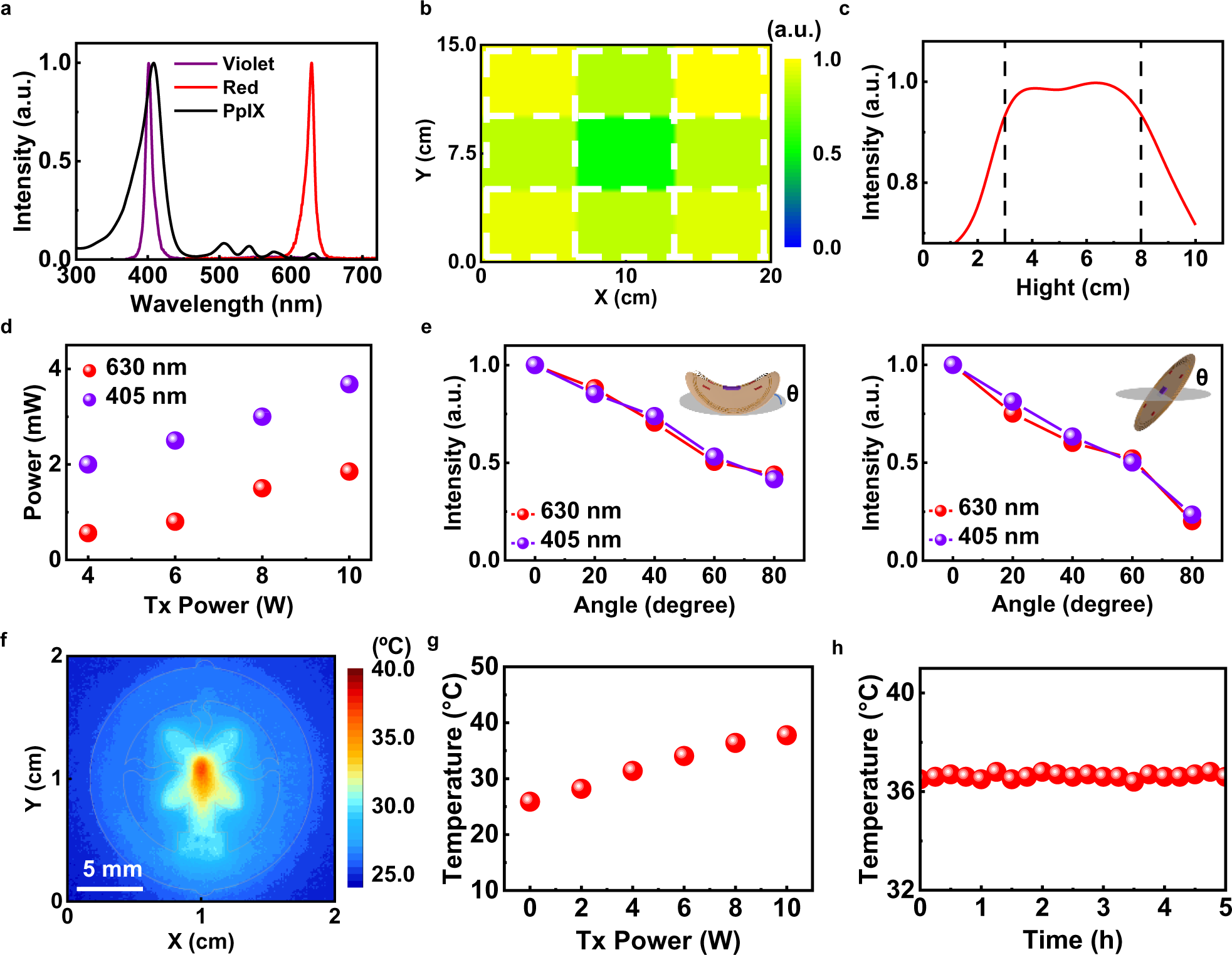
Operational features of the optoelectronic circuit system. (a) Optical spectra for the emission of violet and red microscale LEDs and absorption of PpIX. (b) Light intensity of the coil-powered microscale LEDs placed at different positions on the horizontal plane at a distance of 3 cm from the bottom (c) and on the vertical plane in the center of the cage. (d) Optical power of illuminated violet and red LEDs under different TX power (e) and as a function of bending angle (left) and rotation angle (right), with the circuit placed at a centered position at 3 cm from the bottom of the cage. (f) Measured temperature distribution of the circuit during operation with a TX power of 8 W, a frequency of 50 Hz, and a duty cycle of 65%. (g) Maximum temperature of the operational optoelectronic circuit system as a function of TX power. (h) Maximum temperature measurement of the operational optoelectronic circuit over a period of 5 hours.

Effective thermal management is crucial for active devices that are attached to or implanted in biological tissues. An infrared imager records the temperature distribution across various components of the patch system under varying external RF power levels, including the coil, red and violet LEDs, the rectifier, and capacitors. Figure 2f shows that the most substantial temperature rise occurs at the LED, but it can be controlled by adjusting the duty cycle during pulsed operations(*31*). When subjected to an external RF power of 8 W, with a duty cycle set to 65% and a frequency of 50 Hz, the patch system maintains a maximum temperature below 37 °C. Figures 2g and S5 show that the rise in temperature is influenced by the external RF power, and the modulation of duty cycle and frequency are the most convenient methods for determining the operational temperature of the wireless system. Figure 2h demonstrates the temperature dynamics of the patch over a period of 5 hours, and confirms that the maximum temperature remains within a safe threshold. Within this timeframe, the violet LED emits an energy of 54 J cm^−2^, and the red LED provides 32 J cm^−2^. These energy levels are consistent with the specifications for the light treatment parameters of mPDT.

### Biodegradable MNs

This module comprises densely packed, long pyramidal MNs fabricated from a blend of hyaluronic acid (HA) and polyvinyl alcohol (PVA) affixed to a 0.5 mm thick base layer of PVA. The MNs is a pyramidal shape with dimensions of 700 μm in shaft length, 300 μm in base width, and a tip size of 10 μm. The MNs arrays are fabricated using polydimethylsiloxane (PDMS) negative molds created from cavities shaped like MNs in a metal master mold. The HA-PVA mixture is poured into the PDMS molds to fill the tip and then covered by the pure PVA to form a film. The MNs are dried under vacuum condition and subsequently stored at 4 ℃ for three days to aid in the demolding process, as shown in Figure S6. Figure 3a illustrates the morphology of the biodegradable MNs, which exhibit sufficient flexibility to conform to the curved surfaces of the skin. The MNs are organized into a 40 × 40 array with a 500 μm inter-needle spacing, covering an area of 2 cm ×2 cm. Figure 3b shows that each needle has a height of 700 μm and a base width of 300 μm, achieving a moderate occupancy rate of 60% (the ratio of the cumulative base areas of the MNs to the total area of the array). Considering that the thickness of the skin epidermis varies from 50 μm to 1.5 mm, the length of these needles is sufficient for effective penetration of the skin barrier in applications such as transdermal drug delivery, vaccination, and others, as demonstrated in animal models and human subjects(*32–34*). Figure 3c demonstrates the penetration capabilities of the MNs, as evidenced by the indentations they leave on synthetic skin phantoms covered with a stretched sealing film.

**Figure 3.**
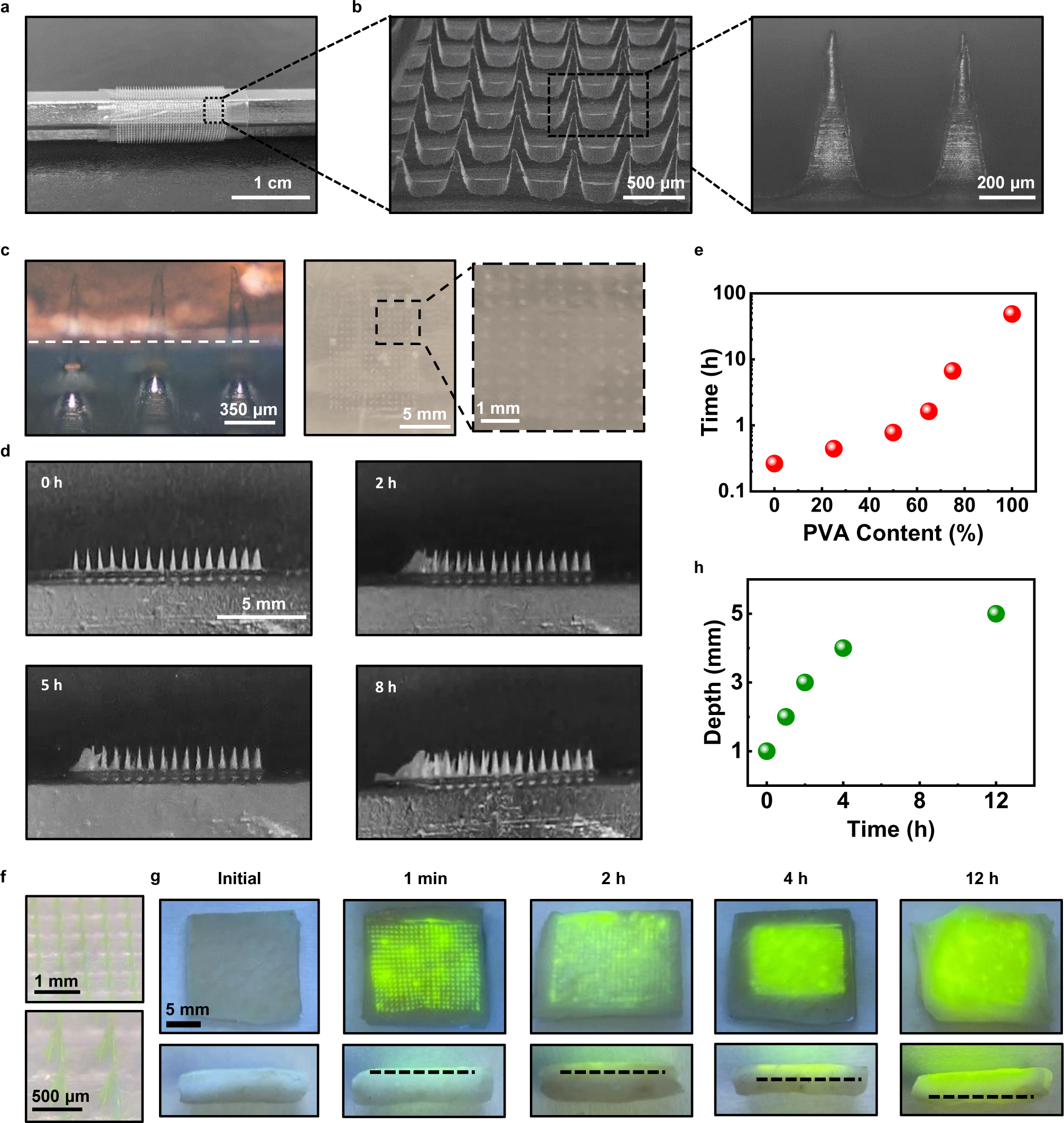
*In vitro* evaluation of degradable MNs. (a) Photographs of the flexible layer of the MNs array applied to the curved surface. (b) SEM images of the MNs. (c) Photographs of an optically mimicking skin phantom covered by the sealing film MNs inserted (left), with imprints remaining on the sealing film after the implantation process (right). (d) Optical images of the MNs applied to the skin phantom for various durations: initial, 2, 5, and 8 hours. (e) The degradation time for different ratios of PVA and HA based on residual MNs morphology. (f) Photographs of MNs with sodium fluorescein tips. (g) Photographs of the diffusion of sodium fluorescein MNs applied to pig skin over time. (h) The diffusion depth of sodium fluorescein as a function of time applied to pig skin.

HA and PVA are chosen for their water solubility, optical clarity, biocompatibility, mechanical resilience, and ease of manufacturing. They also facilitate the efficient embedding of 5-ALA photosensitizers. The proportion of HA to PVA within the MNs is tailored to balance their degradation rates and to ensure adequate rigidity. PVA forms the structural framework of the MNs with its slow dissolution kinetics, crucial for effective tissue penetration and light transmission. Meanwhile, HA is designed to release the encapsulated drug quickly, dissolving within minutes(*35*). The degradation behavior of MNs with varying HA-PVA ratios was systematically studied, with results illustrated in Figures 3d, 3e, and S7. It was observed that the MNs, particularly with a PVA concentration of around 75%, preserved their structural integrity for up to 5 hours. Furthermore, to monitor the distribution of the drug within the MNs and its subsequent diffusion into the skin, a fluorescent dye is utilized as a stand-in for 5-ALA during the production stage, as depicted in Figure 3f. This dye predominantly accumulates at the MNs tips, indicating precise control over its distribution and enabling targeted delivery. Figures 3g and 3h present the diffusion kinetics of the dye release from the degraded MNs with a pig skin model. After applying the MNs to pig skin, sodium fluorescein was observed to gradually permeate the skin layers. This permeation is detectable both at the surface and in cross-sectional views, suggesting that the MNs maintain their transparency and structure while effectively administering the drug into the subcutaneous tissue over a 5-hour therapeutic window.

### Properties of MNs Light Guides

The efficient delivery of UV–visible light to biological tissues is a significant challenge for phototherapeutic methods due to light scattering and attenuation. To demonstrate the efficiency of light transport by the MNs when applied to biological tissues, we have conducted both numerical simulations and experimental characterizations shown in Figure 4. Figures 4a and 4b present the light distribution of UV and red light across various skin depths directly and with the MNs based on Monte Carlo numerical simulations, in which the 2D light intensity profile is normalized to maximum value at the skin surface. The light intensity of violet LED directly applied to the skin decreases to 10% and 1% at propagating distances of 150 μm and 300 μm, respectively. The presence of the MNs enhances the delivery of UV light to the skin at depths of 700 μm and 950 μm, corresponding to the remaining intensity of 10% and 1%. Although the red light owns a relatively good penetration depth, the light intensity attenuates to 10% and 1% at 250 μm and 550 μm with direct illumination and 750 μm and 1200 μm with the MNs, respectively. Therefore, light is redistributed along the insertion depth and transmitted to surrounding tissues through lateral illumination as it passes through the MNs. Compared to direct illumination, the MNs act as light guides and provide efficient UV and red light delivery to the superficial epidermis (up to 100 μm in depth), the papillary dermis (100–300 μm in depth), and the reticular dermis (above 500 μm in depth)(*21, 36*). Numerical simulations show that at UV–visible wavelengths below 500 μm, the optical transmission dose is increased by more than 5-fold (the ratio of optical power after light passes through the MNs to the optical power of the direct illumination), as summarized in Figures 4c and 4d.

**Figure 4.**
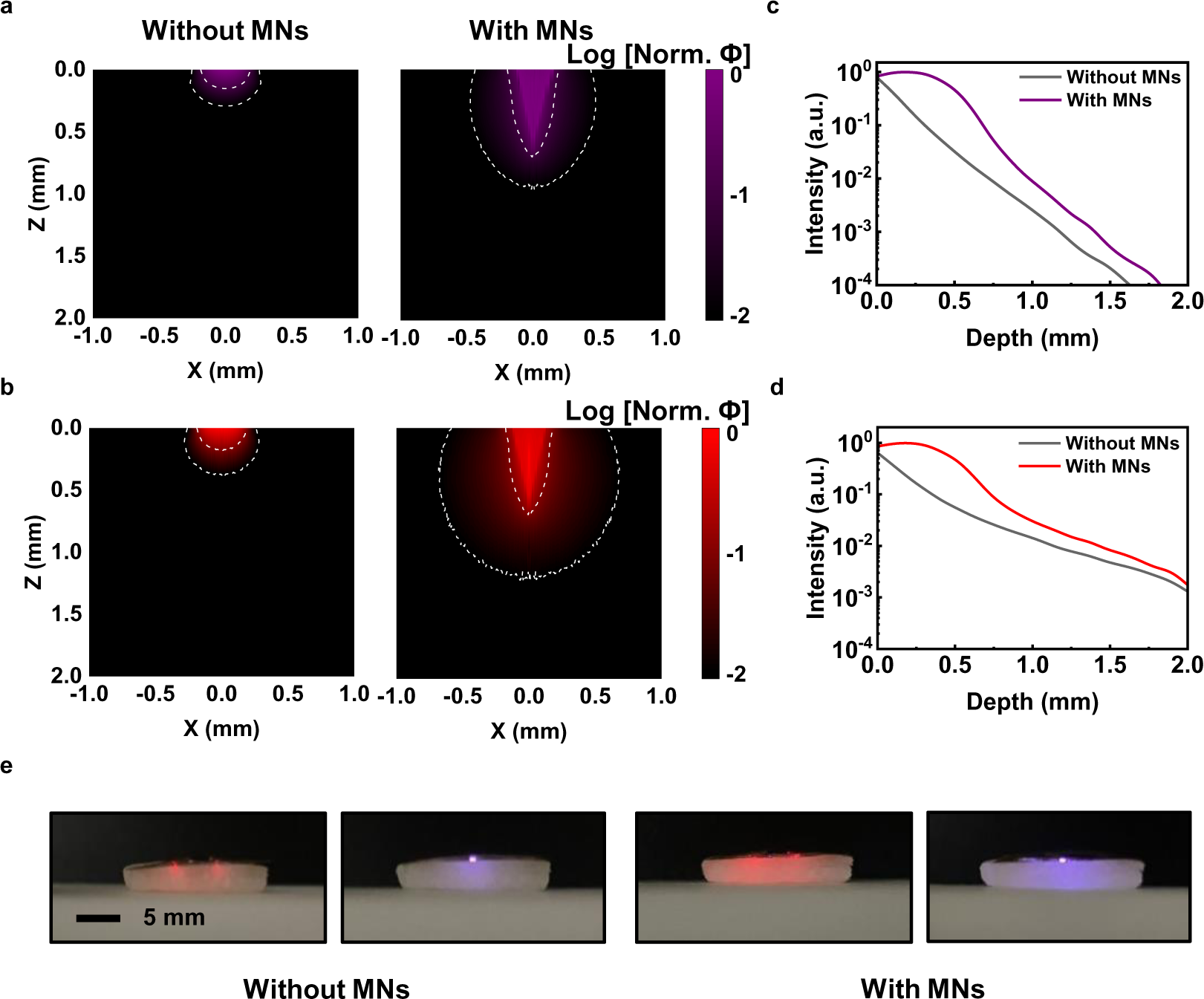
Optical properties of MNs for light delivery. (a) Simulations of optical propagation from the violet LED (b) and red LED under direct illumination (left) and with (right) MNs, which contours depicting the maximum normalized light intensity at 0.1 and 0.01. (c, d) The calculated light intensities from the violet and red LED effectively exposed to skin tissues at various depths under direct illumination and with MNs. (e) Comparison of violet and red light distribution profiles in a skin phantom under direct illumination (left) and in the presence of MNs (right).

Figure 4e shows the experimental results of optoelectric patches irradiating synthetic skin tissue directly or with MNs. The distribution of UV and red light mediated by the MNs covers a larger volume of tissue, which is consistent with the simulation results. This significant difference further demonstrates the importance of MNs in enhancing the penetration of light deep into the tissue. This MNs design opens up new possibilities for treating conditions such as deep-seated localized scleroderma and other skin disorders that have not responded well to conventional phototherapy techniques.

### Assessment in Preclinical Models for Wound Infection

To evaluate the effectiveness and safety of our wireless optoelectronic patch for wound care, we conducted preclinical trials using a mouse model. Figures 5a and S8 show that mice equipped with the wireless optoelectronic patch exhibit comparable mobility to those without the devices. Figure S9 illustrates the use of thermal imaging to continuously monitor the body temperature of mice, confirming that there are no significant temperature fluctuations in the local area. These findings confirm the ability of the wireless optoelectronic patch to maintain comfort without hindering normal behaviors.

**Figure 5.**
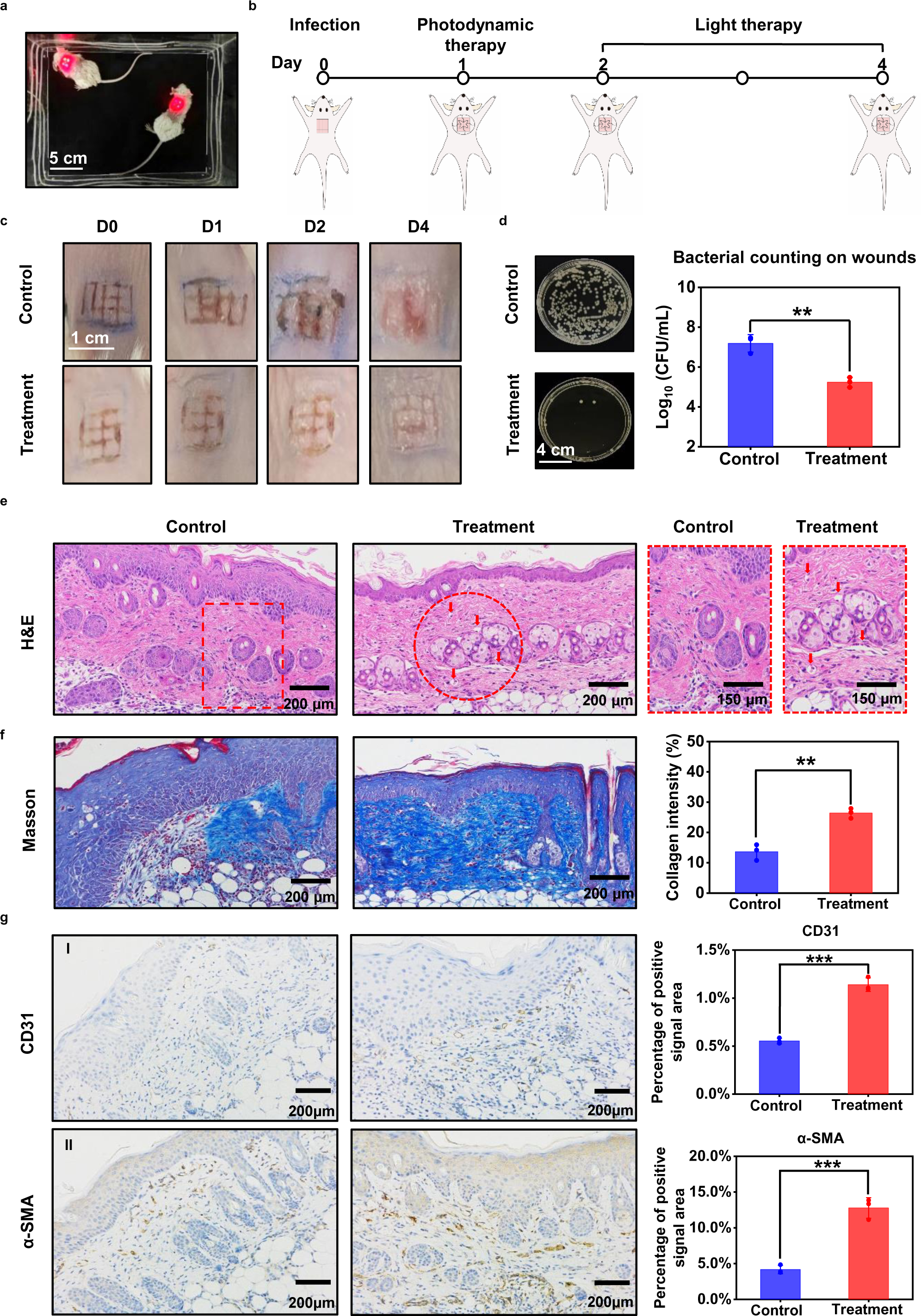
*In vivo* anti-infective therapy with a wearable optoelectronic patch on a wound infected with Pseudomonas aeruginosa. (a) Photograph of freely moving mice wearing optoelectronic patches, with the illuminated LED powered by the wireless circuit. (b) Schematic diagram of the establishment of a mice wound model and treatment processes. (c) Representative optical images of wound healing process at various time intervals (0, 1, 2, and 4 days) after combined treatments of photodynamic therapy and light therapy. (d) Quantitative analysis of bacterial colonies formed by P. aeruginosa collected from different wound tissues. (e) Hematoxylin and Eosin (H&E) staining of skin tissue after 4 days in the control and treatment groups. Arrowheads indicate neovasculature and rectangles indicate a large infiltration of inflammatory cells, highlighted in the right insets. (f) Masson trichrome staining and quantitative analysis of collagen fiber strength in skin tissue after 4 days in the control and treatment groups. (g) Representative images and quantitative analysis of immunohistochemical staining of CD31 (I) and α-SMA (II) in skin tissue after 4 days in the control and treatment groups. All data were presented as mean ± S.D with a two-tailed t-test, ***p < 0.001, **p < 0.01.

In conjunction with comfort evaluations, a series of light therapy sessions are conducted with an optoelectronic patch on the infected wound mice. *Pseudomonas aeruginosa* (*P. aeruginosa*), a Gram-negative bacillus, is well-known for causing opportunistic infections in immunocompromised individuals(*37*). The pathogen is associated with a wide range of clinical conditions, ranging from localized skin problems to serious systemic illnesses. Therefore, the management of *P. aeruginosa* infections related to the skin is crucial, particularly in the context of wound care. Mice in both control and experimental groups are infected with *P. aeruginosa* on 1 × 1 cm² skin wounds, and all subjects develop noticeable inflammation at the site of infection. After inducing a wound infection in mouse skin, a series of interventions are applied with mPDT for its antimicrobial properties, subsequently with multi-wavelength light therapy to reduce inflammation and promote healing, as shown in Figure 5b. Mice in the control group only received conventional sterile wound dressings and were not treated with phototherapeutic interventions. Mice in the treatment group are treated with 5 hours of mPDT on the first day using an optoelectronic patch and MNs loaded with 5-ALA. From the 2^nd^ day to the 4^th^ day, 2 hours of light therapy are performed with the optoelectronic patch. Figures 5c and S10 demonstrate nearly complete wound closure and regeneration of epidermal tissue in the treatment group, while the control group continued to exhibit persistent red and swollen wound symptoms. Figure 5d illustrates the bactericidal impact of the treatment regimen on *P. aeruginosa* within wound sites. The quantitative data show that the groups subjected to the mPDT treatments experienced a significant bacterial reduction, with a 2.48 log_10_ CFU mL^−1^ decrease in *P. aeruginosa* compared to the untreated control group (p < 0.01).

Histopathological examinations are performed to further evaluate the therapeutic effect on the wound skin. Figure 5e presents hematoxylin and eosin (H&E) staining of the wound skin from the control and treatment groups, with the right images providing the details in a magnified view of representative areas. The decrease in inflammatory cells in the treatment group indicates a reduction in bacterial infections of the skin tissue. Meanwhile, the increased neovascularization in the wound area marked by the red arrowheads demonstrates the success of tissue repair and regeneration after phototherapeutic interventions(*38*). Figure 5f presents the Masson trichrome staining and quantitative results of the wound tissue, in which collagen intensity is an essential indicator in the tissue repair process. These results further demonstrate the significantly enhanced strength of collagen fibers in the treatment group compared to the control group (p < 0.01). As the enhanced formation of new blood vessels accompanies changes observed in immunohistochemical assays, the increased expression levels of both endothelial marker CD31 and α-smooth muscle actin (α-SMA) are measured in the treatment group, as shown in Figure 5g, indicating angiogenesis and vascular maturation, respectively. These findings collectively suggest that combination treatment based on metronomic photodynamic and light therapies with the wireless optoelectronic patch is a promising approach for reducing bacterial proliferation and accelerating skin wound healing.

## Conclusion

In this study, we present a wireless optoelectronic patch system for wound care, combining a biodegradable MNs patch with multiple colors of LEDs, which are powered wirelessly via radiofrequency technology. This design eliminates the need for optical fibers or external fixed, rigid light sources, addressing the issues related to weight and battery replacement and enhancing convenience for optical control applications. The biodegradable composite PVA-HA MNs are engineered with an adjustable degradation timeline modulated by the PVA to HA ratio, to allow for tailored therapeutic regimens. These MNs can load the photosensitizer of 5-ALA, ensuring the targeted release of the therapeutic agent through subsequent HA degradation after skin penetration. The residual PVA structure acts as an optical waveguide in the skin, enhancing the spread of excitation light within the tissue. In a mouse model with bacterially infected wounds, the optoelectronic patch successfully administers mPDT in conjunction with complementary light therapy. This combination treatment not only sterilizes wounds but also accelerates healing, demonstrating its potential as an effective and safe solution for wound management, with promise for future clinical adoption.

## Methods

### Fabrication of Flexible Printed Circuit Board

A flexible substrate composed of a copper–PI–copper laminate (with thicknesses of 18/75/18 μm, respectively; DuPont, Pyralux) was subjected to patterned laser ablation using the ProtoLaser U4 (LPKF Laser & Electronics). The circuit interconnects, bonding pads for electronic components, and the geometry, culminating in the formation of the FPCB were precisely defined. To establish electrical connectivity between the top and bottom circuits of the FPCB, laser-plated via holes were filled with conductive silver paste (CD03, Conduction) and subsequently heated to 100 ℃. The components included a surface-mount capacitor (0402CG300J500NT, FH), a rectifier diode (RB161QS-40T18R, ROHM), four red LEDs (RF09E-FB, HC SemiTek Corporation), and a purple LED (URMF23BAS, HC SemiTek Corporation) were hot-air soldered onto the FPCB using solder paste (AZ100, Anderson). To protect and encapsulate the devices integrated into the FPCB, a conformal coating of parylene (12 μm) was applied via chemical vapor deposition. Additionally, a secondary encapsulation layer was introduced using polydimethylsiloxane (PDMS, Sylgard 184, Dow) and subsequently cured in an oven at 70 ℃ for 2 hours.

### Fabrication of Biodegradable Polymer Microneedles

#### Hyaluronic Acid (HA) and Polyvinyl Alcohol (PVA) Solution Preparation

50 mg mL^-1^ hyaluronic acid (HA) solution was formulated by dissolving 5 g of HA in 100 mL of deionized water. Similarly, 50 mg mL^-1^ polyvinyl alcohol (PVA) solution was obtained by heating 5 g of PVA in 100 mL of deionized water on a magnetic heating stirrer at 120°C until fully dissolved.

#### Mixture Preparation

Solutions were formulated with varying HA to PVA ratios, ranging from 1:1 to 1:10, to examine the rate of degradation. A mixture recipe with a 3:1 ratio, consisting of 0.3 g of the PVA solution and 0.1 g of HA solution, was selected for the experiments. Subsequently, 400 mg of 5-aminolevulinic acid (5-ALA) photosensitizer drug (A107209-5 g, Aladdin) was added to these mixtures and stirred until fully dissolved.

#### MNs Casting

The prepared mixed solution was uniformly poured into a PDMS mold, sized 2.5 cm × 2.5 cm, with an effective MNs area of 2 cm × 2 cm. To ensure complete filling of the cavities, the mold containing the mixture was placed in a vacuum drying oven at 24 °C for 20 min. The excess mixture atop the base was meticulously removed, ensuring the solution was confined to the MNs tips. Subsequently, a base layer with a thickness of 500 μm was established by casting 1.5 g of the PVA solution.

#### Drying and Separation

The mold was refrigerated at 4 °C for a drying period of three days, with desiccant beads added to ensure thorough drying. After this period, the MNs patches were gently extracted from the mold, yielding drug-imbued patches of 2 cm × 2 cm dimensions, featuring a 40 × 40 MNs array. These patches were then partitioned into four equal segments, each of size 1 cm × 1 cm, and stored at –20 °C for preservation.

#### Synthetic Skin Phantom Preparation

To demonstrate the light propagation behaviors with or without a MNs in biological scenarios, the skin phantom was prepared to mimic the absorption and scattering properties of living tissues.(*39*) In brief, the phantom was composed of agarose (A801450-0.6 g, Macklin), intralipid (3.0 g, 20% emulsion, Sigma Aldrich), all fully dissolved in a phosphate buffer solution with a pH of 7.4. The mixture is boiled, allowed to cool naturally to room temperature, and then solidifies into a gel-like form.

### Device Characterization and Modeling

#### Optoelectronic Characterizations

The emission spectra of the violet and red LEDs were measured using a spectrometer (HR2000+, Ocean Optics). The current–voltage characteristics of the LEDs were recorded by Keithley 2400. The emission intensity of the LEDs was recorded with a calibrated silicon photodiode (SG2387-9898E, SGECL). The temperature distributions on the circuit board are mapped with an infrared camera (FOTRIC 228).

#### In Vitro Release Assays

The drug was loaded into the MNs with fluorescein sodium (F809553-25g, Macklin) instead of 5-ALA, allowing for clear visualization of the delivery, as shown in Figure 3g. The MNs patch was applied to pig skin. Both the surface and cross-section of the skin were examined for the percentage of cumulative drug release and the dynamic diffusion depth. All observations were documented under the illumination of a 15 W 365 nm UV lamp using a microscope (XTL-165-MT).

#### Optical Modeling

Monte Carlo simulations were performed using the TracePro Free Trial Version to examine light propagation into skin tissue and derive related curves. Two simulation scenarios were evaluated to assess the light distribution in the tissue under a 300 µm × 300 µm light source with Lambertian radiation at wavelengths of 405 nm and 630 nm, which are examined both directly and with MNs. The parameters of skin tissue to purple light and red light in the simulation included scattering coefficient of 26.9 mm^−1^ and 33.84 mm^−1^, absorption coefficient of 0.75 mm^−1^ and 0.068 mm^−1^, anisotropy factor of 0.78 and 0.874, and refractive index of 1.36(*40*). The MNs were set to a bottom diameter of 300 µm, a top diameter of 10 µm, a height of 700 µm, a refractive index of 1.5, a transmittance of 1, and an absorption coefficient of 0.

### Biological Studies

#### Animals

All animal procedures were approved by the Ethics Committee of Chinese PLA General Hospital. Balb/c female mice purchased from the Vital River Laboratory (Animal Technology, Beijing, China), aged 6–8 weeks and weighing between 18–20 g, were individually housed in a sterile environment. Animals were maintained on a 12 h light/dark cycle at 22–25 °C and provided with access to food and water ad libitum.

#### Bacteria Infection

*Pseudomonas aeruginosa* (*P. aeruginosa*, ATCC 28357) was sourced from the Department of Microbiology at the Chinese People’s Liberation Army General Hospital in Beijing, China. Single colonies of *P. aeruginosa* were inoculated in tryptic soy broth (TSB, Oxoid, UK) with 1% glucose and incubated overnight at 37℃ with agitation at 200 rpm. Based on measurements taken with DENSIMAT (BioMérieux), the bacterial suspensions were then diluted to a 0.5 MCF standard (equivalent to 1.5×10^8^ CFU/mL) in TSB. Twelve healthy mice being anesthetized with isoflurane (Reward, Shenzhen) and having their fur shaved, 1 × 1 cm^2^ skin wounds marked by 6 × 6 crossed scratch lines were created on the dorsal surface of the mice using 26-gauge needles(*41*). Each wound was inoculated with a 50 µL suspension of *P. aeruginosa* at a concentration of 1.5 × 10^8^ CFU mL^−1^, allowing the bacterial suspension to be absorbed into the wound.

#### Metronomic Photodynamic Therapy and Light therapy

Twelve mice were randomly divided into control and treatment groups, with six mice in each. Mice in the control group had their wounds covered with sterile gauze and were subsequently placed in sterile cages without further intervention. Conversely, mice in the treatment group underwent mPDT using degradable MNs infused with 5-ALA, followed by light therapy. On the first day, a biodegradable MNs patch loaded with 5-ALA was applied to the wound and left for an incubation period of 30 minutes. Afterwards, the mice in the treatment group underwent PDT for a duration of 5 hours using coil-powered violet and red LEDs. In the days following, the mice in the treatment group received daily 2 hours sessions of light therapy. These violet and red LEDs operate at powers of 3 mW and 1.8 mW, respectively, and both maintain a 65% duty cycle and a frequency of 50 Hz.

#### Histopathological Examination

Upon concluding PDT and light therapy on three mice from each treatment group, a 1 cm^2^ section of the wounded skin was extracted from each euthanized mouse. These tissues were fixed overnight in 4% paraformaldehyde, then sequentially dehydrated using ethanol and embedded in paraffin. After being sectioned into slices of 3 – 5 µm thickness, the slices were stained with hematoxylin-eosin (H&E), Masson trichrome staining CD31 antibody, and α-SMA antibody to assess histological alterations and evaluate the inflammatory response, respectively.

Histological and immunohistochemistry images were captured using an optical microscope (Nikon, Eclipse CI) and recorded with a digital camera (Nikon, DS-U3). For image analysis, ImageJ software was employed to binarize images captured under consistent settings. Staining quantification was determined by calculating the percentage of positively stained area to total area based on a set intensity threshold. A two-tailed paired t-test was utilized to determine statistical significance, with a p-value of < 0.05 indicating a statistically significant difference.

## Acknowledgments

This work is supported by the Beijing Nova Program (20230484254), the National Natural Science Foundation of China (NSFC) (62005016, 62025104, T2293750, T2293753), the National Key Research and Development Program of China (2022YFA1207600).

## Author contributions

Z. X., X. R., and H. D. performed device design, fabrication, and characterization. Z. X., W. T., X. R., Y. X., and H. D. designed and tested circuits. Z. X., W. C., T. S., Y. X., Z. C., Y. G., H. Z, and H. D. designed and performed biological experiments. X. S., Y. W., H. Z., J. Y., and H. D. provided tools and supervised the research. Z. X., W. C., H. Z., J. Y., and H. D. wrote the paper in consultation with other authors.

---

**Figure S1.**
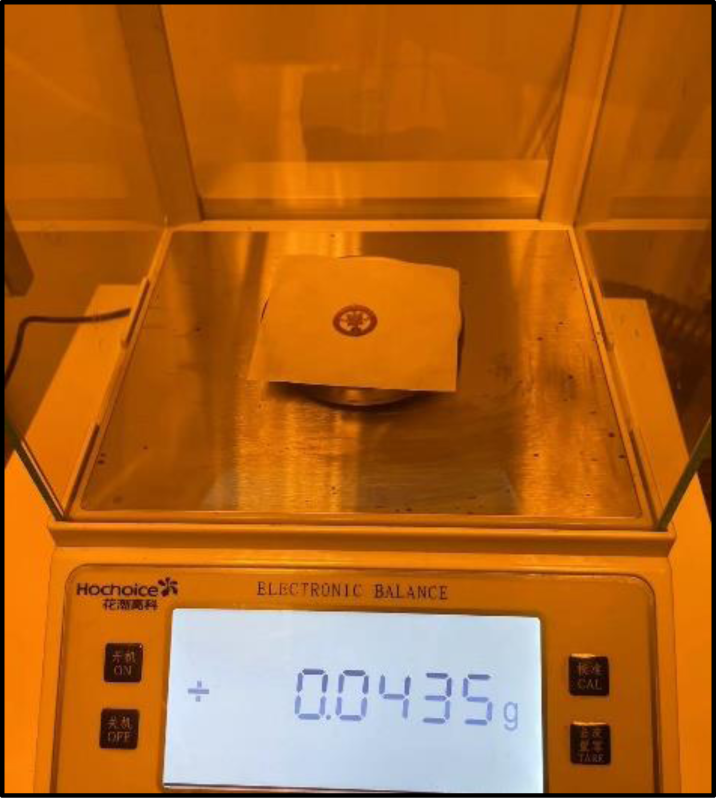
The weight of the battery-free optoelectronic patch.

**Figure S2.**
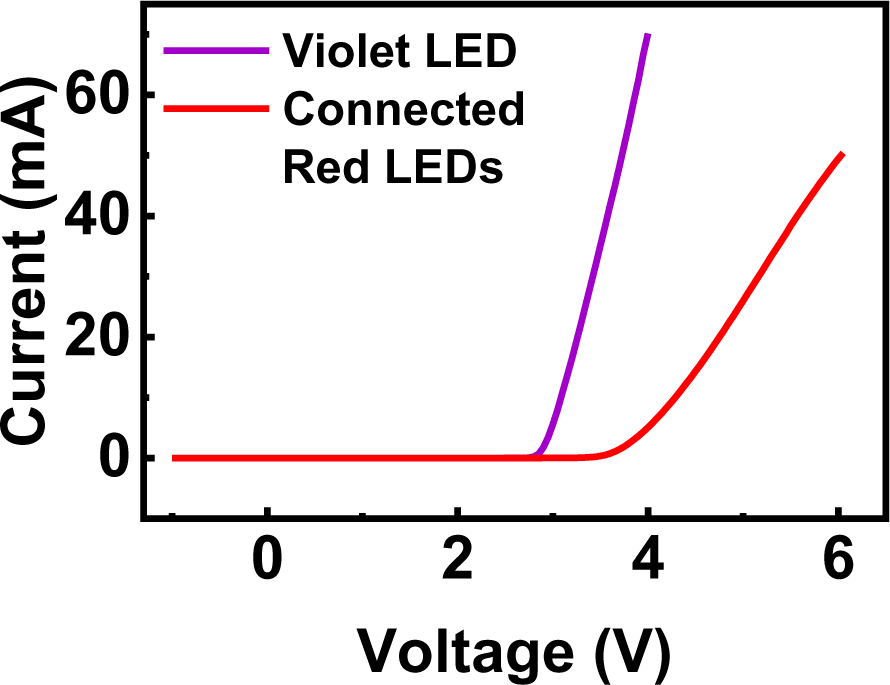
The current – voltage characteristics of a violet LED and 4 red LEDs connected in 2 series and 2 parallel configurations.

**Figure S3.**
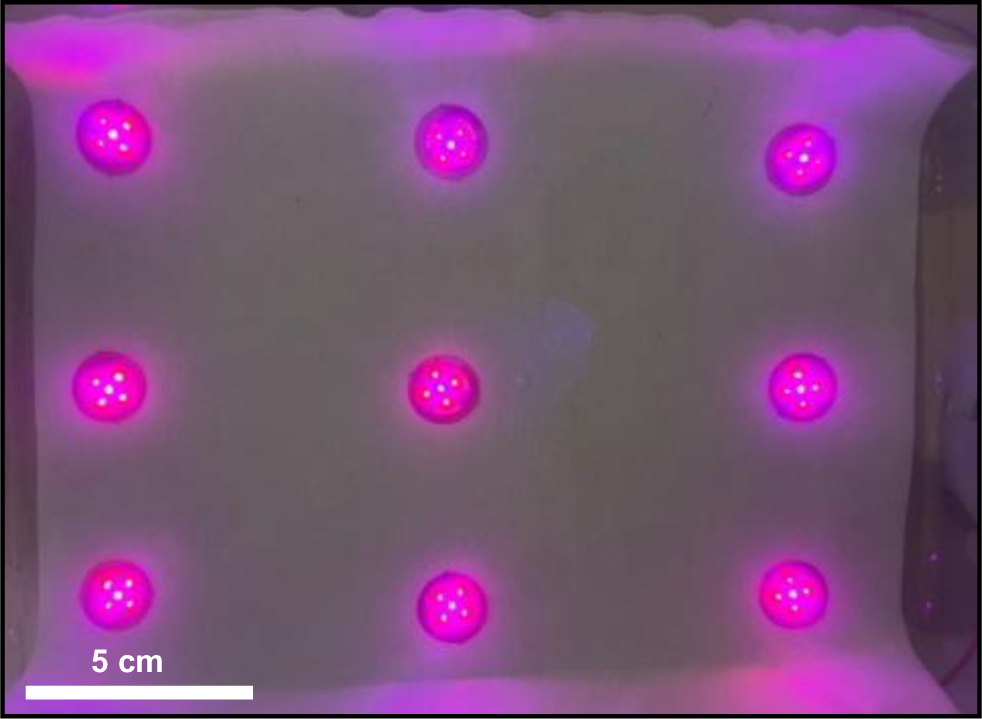
Wireless operation of 9 devices placed in the home cage (20 cm × 15 cm) at a height of 3 cm from the bottom.

**Figure S4.**
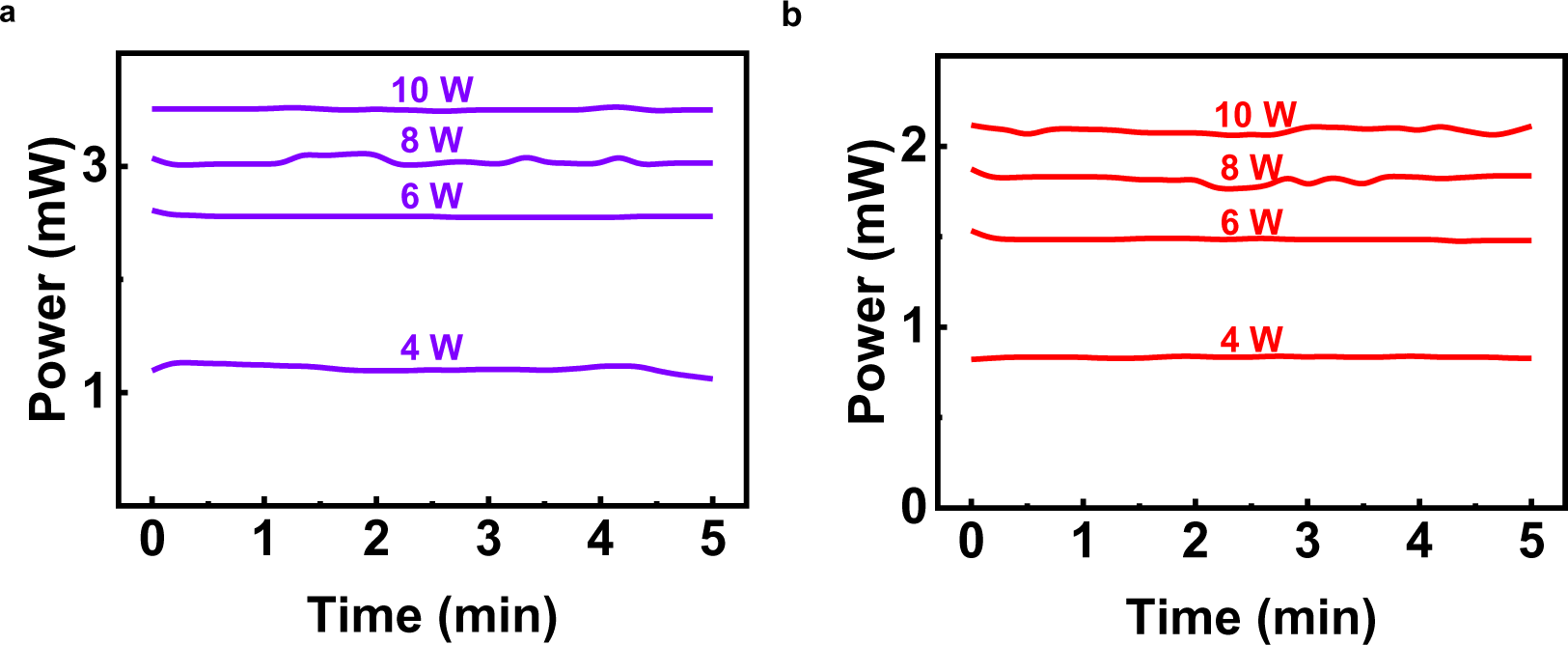
The light optical power of coil-powered (a) a violet LED (b) and 4 red LEDs under various TX powers.

**Figure S5.**
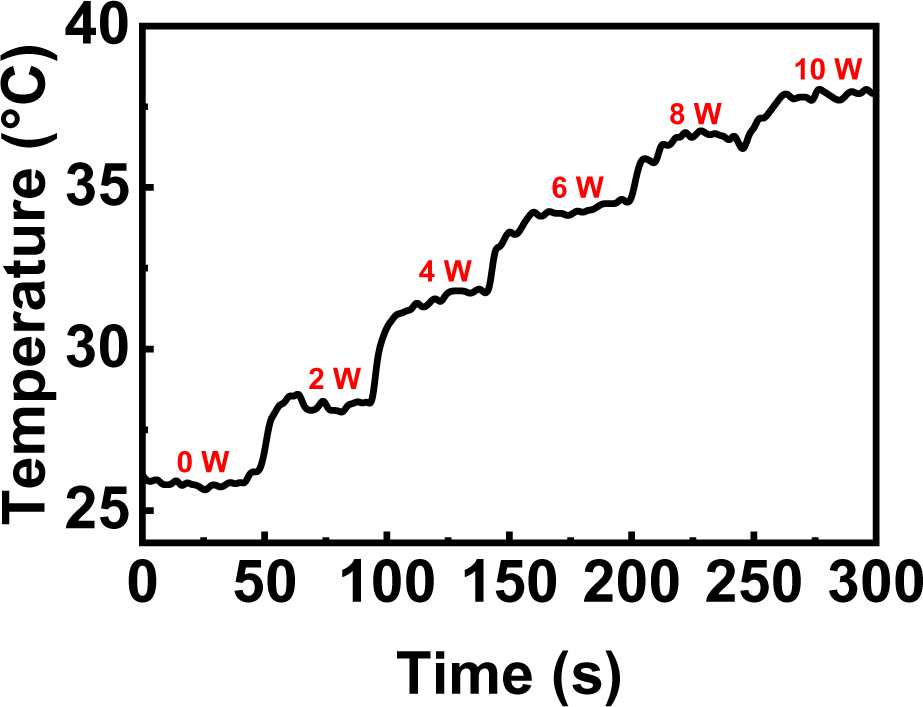
The maximum temperature changes for the operational optoelectronic circuit under step-tuned TX power from 0 W to 10 W.

**Figure S6.**
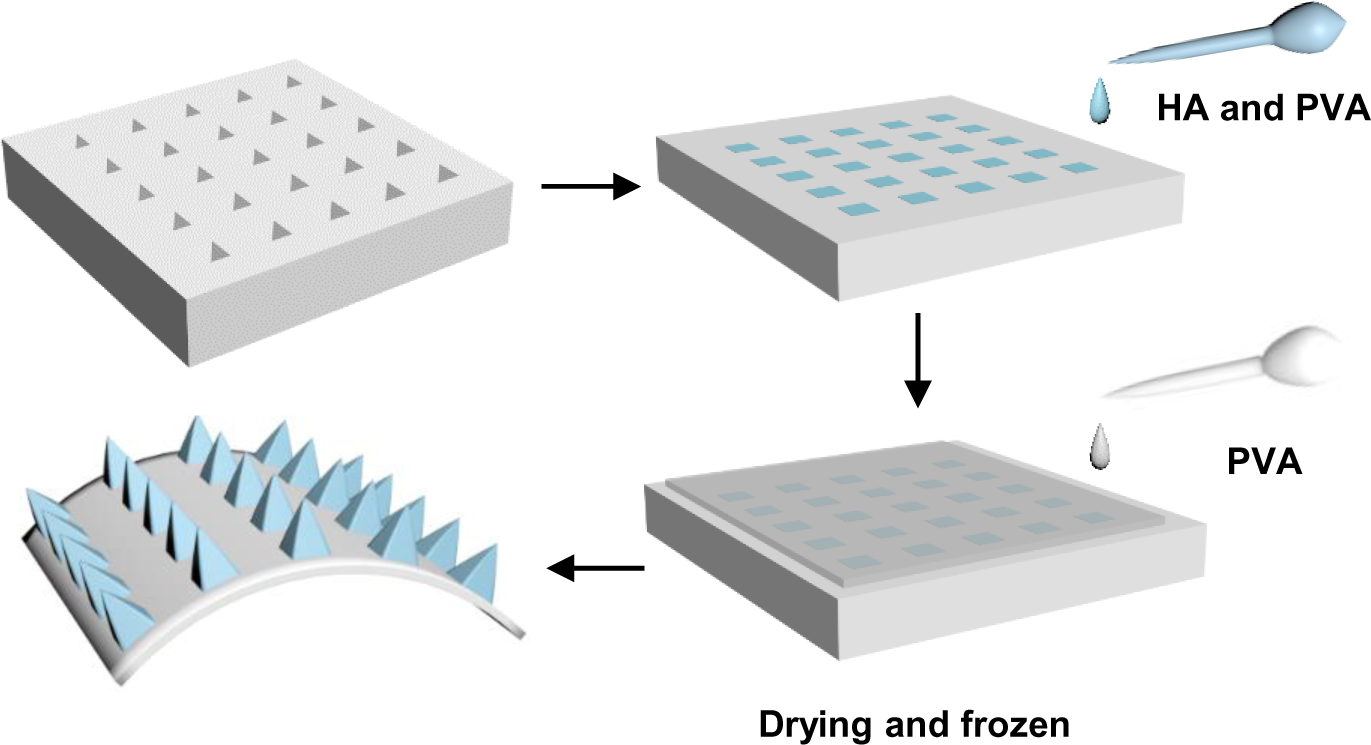
Schematic illustration of the fabrication process for the biodegradable MNs.

**Figure S7.**
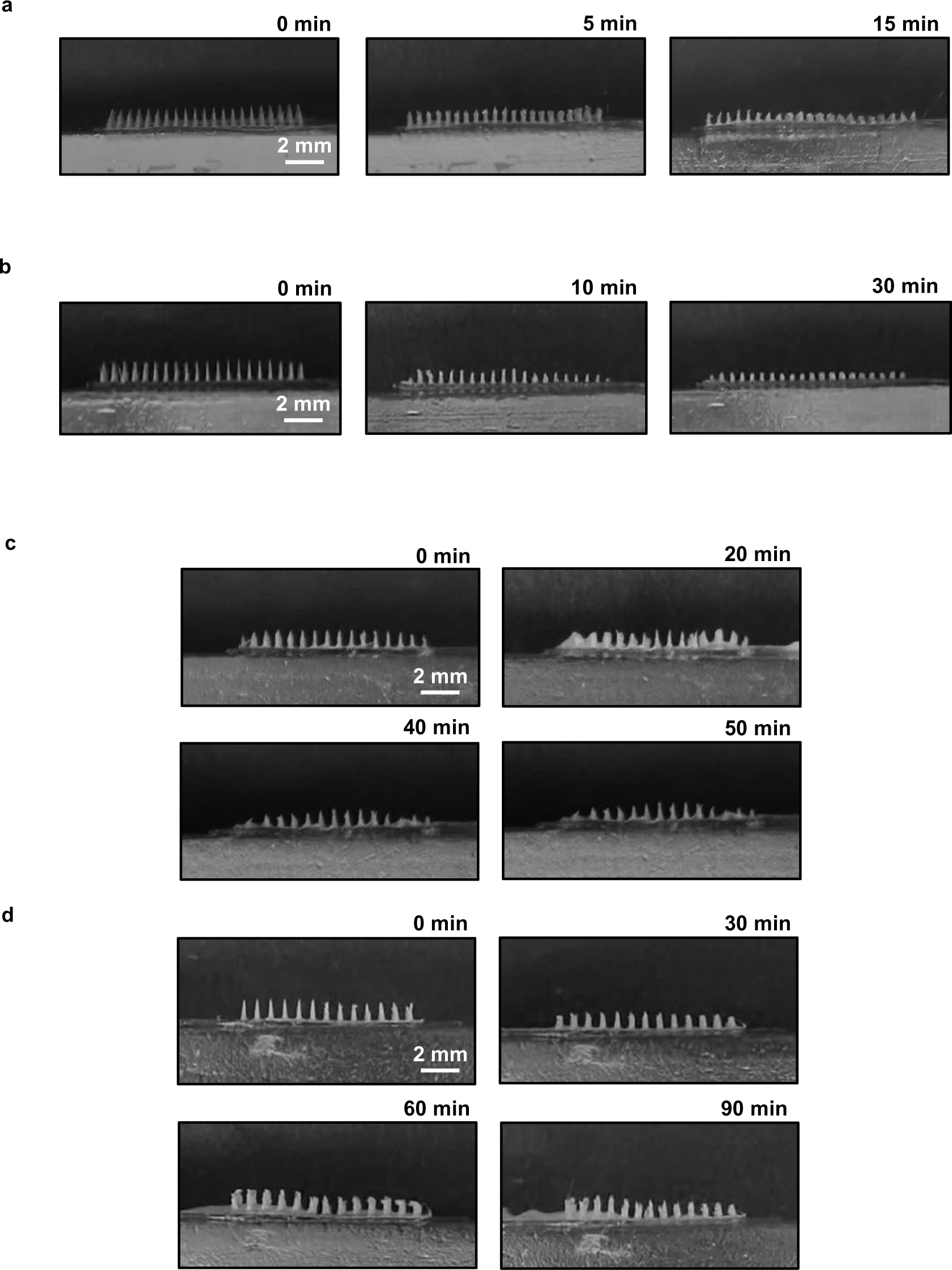
Optical images of the MNs applied to the skin phantom at various times with the proportion of (a) 0%, (b) 25%, (c) 50%, and (d) 65% PVA.

**Figure S8.**
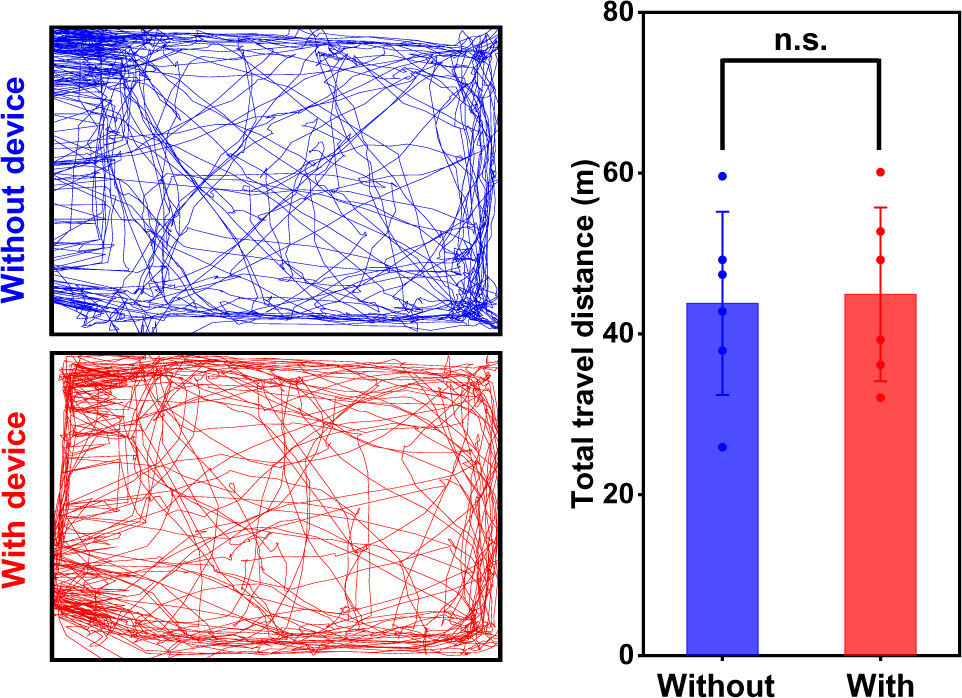
Mobility comparisons between mice moving freely and with optoelectronic patch in the open-field test (left), and statistical analysis between the two groups (right). 6 mice are used in each group (n.s. P > 0.05).

**Figure S9.**
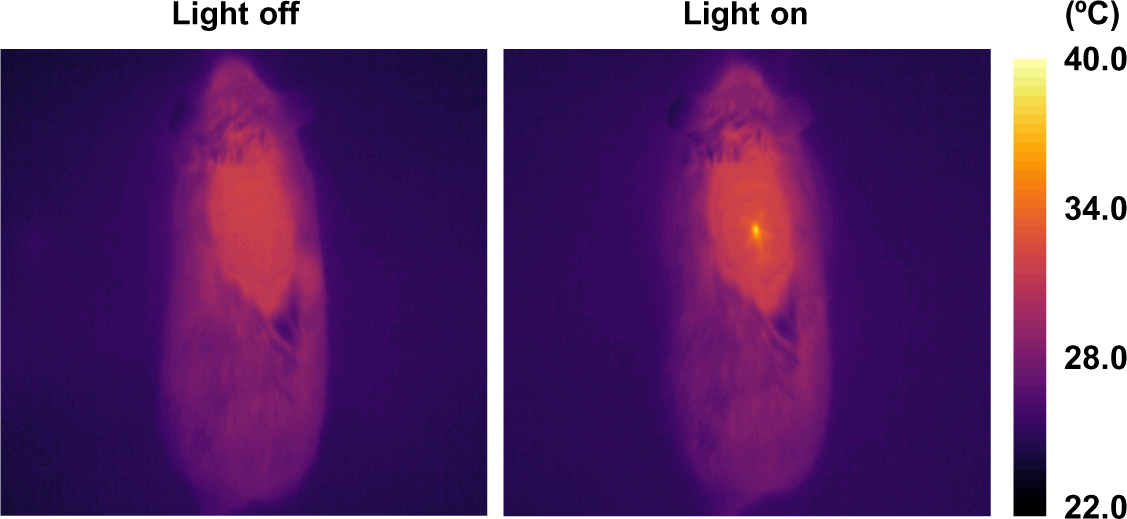
Heat maps of mice with optoelectronic patches in non-operational (light off) and operational (light on) states.

**Figure S10.**
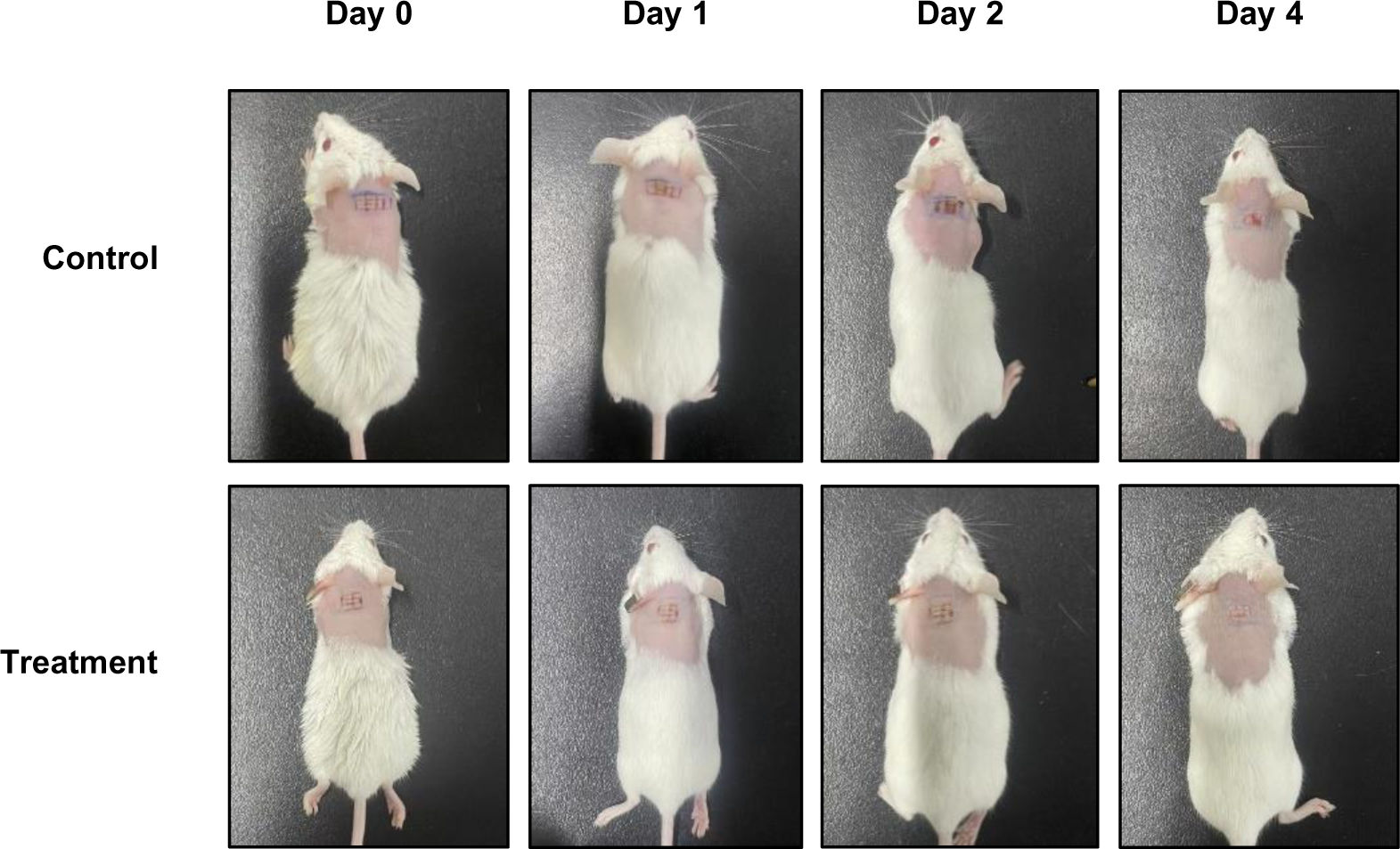
Optical images of the mice during the wound healing process at various time intervals (0, 1, 2, and 4 days) after combined treatments of photodynamic therapy and light therapy.

